# Challenging the impact of consortium diversity on bioaugmentation efficiency and native bacterial community structure in a freshly PAH-contaminated soil

**DOI:** 10.1101/2024.04.13.589391

**Authors:** E.E. Nieto, S. Festa, D. Colman, I. S. Morelli, B. M. Coppotelli

**Affiliations:** Centro de Investigación y Desarrollo en Fermentaciones Industriales, CINDEFI (UNLP; CCT-La Plata, CONICET), La Plata, Argentina; Comisión de Investigaciones Científicas de la Provincia de Buenos Aires, La Plata, Argentina

**Keywords:** bacterial consortia, allochthonous bioaugmentation, ANCOM-BC, substrate availability, inoculum establishment

## Abstract

Polycyclic aromatic hydrocarbons (PAHs) are priority pollutants. We studied the effect of bioaugmentation with three allochthonous bacterial consortia with increasing diversity, SC AMBk, SC1 and SC4, in the structure and functionality of an acutely PAH-contaminated soil microbiome. The PAH supplementation increased the resource availability and the inocula were able to: efficiently degrade the PAHs supplemented after 15 days of incubation, become temporary established, and modify the number of total interactions with soil residents. *Sphingobium* and *Burkholderia,* both member of inoculants, were the major contributors to KO linked to degradation and to differentially abundant genera in inoculated microcosms, indicating their competitiveness in the soil. Bioaugmentation efficiency relayed on them, while further degradation, could be carried out by native microorganism. This is the one of the first works which applied three inocula, designed from naturally occurring bacteria and study their effect on the soil native community through the ANCOM-BC. We revealed that when a resource that can be use by the inoculant is added to the soil, it is not necessary a high-diversity inoculant to interact with native community and establish itself. This result has implications in the design of microbiome engineering for bioremediation processes

## 1. Introduction

Polycyclic aromatic hydrocarbons (PAHs), produced mainly by anthropogenic sources, are priority organic pollutants, according to the US Environmental Protection Agency, affecting the health of humans and other organisms (Ravindra et al., 2008; García-Sánchez et al., 2018; Patel et al., 2020). The metabolism of some PAHs generates intermediates (e.g., diol epoxides, quinones, hydroxyalkyl derivatives), which form covalent adducts with nucleic acid and lead to genotoxic effects (Abdel-Shafy and Mansour, 2016). Due to their hydrophobicity, these contaminants are easily adsorbed into soil particles and, therefore, soils function as a reservoir of PAHs (Kuppusamy et al., 2017).

Bioremediation is a cost-effective and ecofriendly technology to reduce the concentration of pollutants from environmental matrices (Patel et al., 2020). In order to do so, it is key to apply in situ microbiome engineering approach, which aims to control and design community-level properties through (Lawson, 2021) the manipulation of indigenous microbes or the introduction of new members (Albright et al., 2022). As microbiome engineering through bioaugmentation, seeks to generate microbiomes with new functionalities (e.g. degradation of a target pollutant), it has been proposed as a powerful tool to improve environmental bioremediation of PAH (Wanapaisan et al., 2018; Laothamteep et al., 2021).

An effective strategy for bioremediation of acutely contaminated soils is the inoculation of allochthonous PAH-degrading microorganisms (Nwankwegu et al., 2022). The use of microbial consortia as inocula in bioremediation processes became relevant in recent times, since the interactions between their members can optimize the degradation processes (Massot et al., 2022). Positive interactions favor consortia stability and robustness, which can be correlated with consortia diversity, and increase the probability of inoculum establishment (Stenuit and Agathos, 2015; Che and Men, 2019; McCarty and Ledesma-Amaro, 2019; Ziesack et al., 2019). Also, pre-existing interactions in consortia can be exploited by creating highly specialized microbial units where organisms fulfill certain niches (Albright et al., 2022).

A crucial aspect in bioremediation processes is the understanding of microbial diversity, community structure, interactions, and metabolic activities (Muangchinda et al., 2018). Soil microbes co-exist in complex networks with multiple interactions that influence the functional stability of soil ecological system(Fath et al., 2007; Wang et al., 2018; Shi et al., 2020; Tu et al., 2020). A novel perspective posits that microbial communities are organized into metabolically cohesive units where consortia with positive feedback loops use resources in a stable manner and minimize competition through resource specialization and exclusion of generalists (Pascual-García et al., 2020). Synthetic consortium has been described as forming stable microbiomes with previously established community members, as long as key players can compete with resident microorganisms (Lawson, 2021).

Bioaugmentation strategies could fail due to the lack of microorganisms performing critical functions (Iwabuchi et al., 2002), or because consortia selected under laboratory conditions are outcompeted by the native community in the field (Palkova, 2004; Steensels et al., 2019). Monitoring inoculants is needed to determine whether the desired functional changes were achieved as a consequence of the establishment or modification of desired functions in the community (Cornell et al., 2021; Albright et al., 2022). Furthermore, the introduction of new species during bioaugmentation can have a relevant impact on the native community, even when the abundance of the inoculant decreases (Mallon et al., 2018; Mawarda et al., 2020). Therefore, soil microbiome engineering efforts should consider the assessment of establishment of the inoculants and the footprint left in the native community after the bioaugmentation process and not only inoculant characteristics (Kaminsky et al., 2019; Manfredini et al., 2021).

In previous works, three PAH-degrading synthetic consortia from naturally occurring bacteria were designed: SC AMBk, SC1 and SC4 consortia, which differed in number and composition of bacterial species (Macchi et al., 2021; Nieto et al., 2023). Their degradation potential was evaluated encompassing the individual strain roles, the interaction among strains and time-driven changes in gene expression and metaproteomic patterns during PAH removal. These consortia increased, due to synergistic interactions, the degradation efficiency of PAHs in liquid medium in relation to axenic cultures, positing them as good candidates for application in bioaugmentation processes (Macchi et al., 2021; Nieto et al., 2023).

This work evaluates the effects of the three allochthonous bacterial inocula, SC AMBk, SC1 and SC4, in the structure and functionality of a native microbial community of an acutely contaminated soil and monitors the constituent strains in soil during the bioaugmentation process. We hypothesized that when using a more diverse inoculant in bioaugmentation strategies, an improvement in pollutant degradation activity will be observed due to different microbiome assemblies. This was carried out as part of a continued effort to develop and refine an *in situ* microbiome engineering scheme to remediate PAH-contaminated sites. The novelty of this work is the use of three inocula designed from naturally occurring bacteria for the bioaugmentation experiments, the monitoring of the constituent strains of the inoculated consortia and the comparison of the effect on the soil native community through the ANCOM-BC (Analysis of Compositions of Microbiomes with Bias Correction).

## 2 Materials and methods

### 2.1 Soil Samples

Non-contaminated soil (SP) samples were collected from an urban park near La Plata city, Argentina (34°51’24.6“S;58°06’54.2”W). Physicochemical properties were determined in the Laboratory of Soil Science at the National University of La Plata. SP soil was a clay loam soil with a pH of 5.8-5.9, 3.60% organic carbon, 6.21% soil organic matter, 0.296% total nitrogen, 4.2 mg.kg^-1^ available phosphorus and no hydrocarbons were detected.

### 2.2 Defined Consortia preparation

SC AMBk, SC1 and SC4 were constructed with different combinations of the strains *Sphingobium* sp AM*, Burkholderia* sp Bk, *Pseudomonas* sp Bc-h and T, *Inquilinus limosus* Inq, and *Klebsiella aerogenes* B, all belonging to an enrichment culture CON (Festa et al., 2013; Macchi et al., 2021). Each strain was grown independently in R3 broth at 28°C and 150 rpm overnight, then the cells were harvested by centrifugation at 6000 rpm and washed twice with sterile NaCl 0.85% (w/v). Cell number of each suspension was determined measuring O.D at 580 nm. SC AMBk was composed of the strains AM and Bk in a 65:35 ratio, being AM the major strain; SC1 was composed of the strains AM, T, Bc-h, B and Inq; and SC4 of AM, Bk, T, Bc-h, B and Inq. Both SC1 and SC4 were built with equal proportions of each strain.

### 2.3 Microcosms set up

Microcosms of SP soil were constructed with 150 g of sieved soil (2 mm mesh) in 350 ml glass containers. SP soil was contaminated with a mix solution containing 150 mg.kg dry soil^-1^ of fluorene (FLU), 600 mg.kg dry soil^-1^ phenanthrene (PHE), 100 mg.kg dry soil^-1^ anthracene (ANT), and 150 mg.kg dry soil^-1^ pyrene (PYR) to a final concentration of 1000 mg.kg dry soil^-1^ of PAH. The spiking solution was added one day before inoculation. Five treatments were carried out in triplicate: 1) microcosm without contamination or inoculation (non-contaminated or control); 2) microcosm contaminated with the spiking solution but without inoculation (SP+PAH); and the three inoculated treatments were inoculated with 5*10^7^ CFU.ml^-1^ of 3) SC AMBk; 4) SC1; 5) SC4. The final moisture was adjusted to 20% humidity. All microcosms were incubated during 58 days at 24±°C and were mixed weekly for aeration.

### 2.4 PAH quantification

PAH concentration was determined on 5 g of soil sampled from each microcosm at 0, 15, 30, and 58 days of incubation. Samples were lyophilized (L-3, REFICOR) and three consecutive extractions were carried out, using 15 ml of hexane:acetone 1:1 (v/v). The hydrocarbons were extracted in an ultrasonic bath (Testlab Ul-trasonic TB10TA) at 40 kHz, 400 W for 30 min. The mixture was centrifuged at 3000 rpm for 10 min and the supernatants were collected in brown glass flasks and evaporated. Then, samples were resuspended in 2 ml of hexane: acetone and filtered (0.45-μm nylon membrane). 5 μl of each sample was injected into a Perkin Elmer Clarus 500 GC-FID. The oven temperature was programmed from 100 °C (initial time 2 min) to 170°C (40 °C.min^-1^) in the first step, 190°C (10°C.min^-1^) in the second step, and to 250°C (40°C.min^-1^), following and extension phase of 2 min. The retention times of the different PAH were determined with standard solution and quantified using calibration curves through serial dilutions.

### 2.5 DNA extraction and Real time PCR

Total DNA was extracted from 1 g of soil from each triplicate microcosm at 0, 15, 30 and 58 days using the e E.Z.N.A. Soil DNA Kit (Omega Bio-Tek, Inc., USA) following the manufacturer’s instructions, and DNA was quantified with NanoDrop 2000 (Thermo Fisher).

Copy number of PAH-degradation related genes: PAHs ring-hydroxylating α subunit-dioxygenase of Gram-negative (GN PAH-RHDα) (Cebron et al., 2008) and catechol 2,3 dioxygenase (C230) (Sei et al 1999), and eubacterial 16S rRNA gene (Harms et al 2003) were quantified by qPCR in a Rotor-Gene Q 2plex (QIAGEN). qPCR reactions were performed in 10 μl final volume, contained 2 μl of template DNA, 5 μl 2x SsoAdvancedTM Universal SYBR Green Supermix (Bio Rad), 1 μl of each primer (0.4 mM) and MiliQ water. Technical triplicates of each biological replicate were performed. The amplifications were carried out with the following cycling parameters: an initial heating to 95 °C (5 min), followed by 40 cycles (16S rRNA) or 50 cycles (PAH-RHDαGN and C203). These cycles consisted of 30s of denaturation at 95 °C, 30s at the primer’s specific annealing temperature (53°C for 16S rRNA and 57°C for GN PAH-RHDα and C230) and 30s of elongation at 72 °C. The final step was for 7 min at 72 °C. Melting curve analyses were carried out to dismiss contamination. Sample gene-copy number was determined using standard curves built from serial dilutions of gene-target specific plasmids.

### 2.6 Sequencing Analysis

Total DNA of day 0 of control microcosms, and day 15 and 58 samples of all treatments were selected to analyze bacterial community composition using V4 hypervariable region of bacterial 16S rRNA gene using universal primer set 515F (GTGCCAGCMGCCGCGGTAA) 806R (GGACTACHVGGGTWTCTAAT). Sample sequencing was performed using Illumina NovaSeq 6000 platform at Novogene Company.

Paired end sequencing data was analyzed using QIIME 2 (v2022.2) (Bolyen et al., 2019). Raw data was filtered followed by denoising with DADA2 to obtain Amplicon Sequence variants (ASV). Taxonomy was assigned using SILVA v.138 trained database.

To follow the behavior of inoculated strains, the ASVs with higher sequence identity with the 16S rRNA gene sequence for each strain were identified using BLASTn database. These ASV were used to calculate their contribution of each strain to its respective genus.

Alpha diversity metrics (Chao1, Shannon and Faith’s Phylogenetic Diversity) were estimated using q2-diversity after samples were rarefied to 72046 reads per sample. Further analyses were carried out in the R environment. Beta diversity was analyzed through a Principal Coordinates Analysis (PCoA) using Bray-Curtis dissimilarity index at ASV level, performed with the *phyloseq* (v.1.42.0) and vegan (v.2.6-4) R packages. PICRUSt2 software was utilized to predict the abundance of gene families KEGG (Kyoto Encyclopedia of Genes and Genomes) orthologs (KOs) (Douglas et al., 2020). Differential abundance analysis of both taxonomy (at genus level) and functions (at KO level) were carried out using ANCOM-BC (v2.0.0) R package (Lin and Peddada, 2020). A prevalence filter of 0.1, a significance of *p<*0.05 and multiple pairwise comparison considering the mdFDR, using holm as adjusted method.

To create the co-occurrence networks for each study, ASVs tables were filtered with relative abundance above 0.5 %. Correlation Network Analysis was performed using Sparse Co-occurrence Network Investigation for Compositional data (SCNIC) integrated in the q2-scnic plugin from Qiime2 software toolkit. From correlations matrix the networks were constructed with a minimum of R correlation-value cutoff of 0.50 (p-value 0.01). For each microbial network, topological parameters were measured by Network Analyzer plugin in Cytoscape v3.7.9. Positively interconnected ASVs were identified and visualized using Gephi v0.10.

### 2.7 Statistical Analysis

Statistical analysis of PAH concentration and diversity indexes were performed through mixed ANOVA. Normality and homogeneity of variances assumptions were checked through Shapiro-Wilk and Levene test, respectively. Pairwise comparisons were corrected using Benjamini-Hochman method. All the tests were performed using *rstatix* (v.0.7.2) R package.

### 2.8 Data deposition

The raw data sequence for 16S rRNA gene amplicons have been deposited under NCBI accession number PRJNA889803.

## 3 Results

### 3.1 Degradation of supplemented PAHs

Degradation efficiency of the bioaugmentation process with the three different consortia was evaluated by analyzing the remaining concentration of the supplemented PAHs (fluorene, phenanthrene, anthracene and pyrene) along incubation time (0, 15, 30 and 58 days) in the microcosms inoculated with SC AMBk, SC1 and SC4 consortia as well as in SP+PAH (contaminated control without inoculation). Results are shown in figure 1. The non-contaminated control (control) was excluded from this figure along with day 58 of all treatments, as PAHs concentration was below the detection limit.

**Figure 1:**
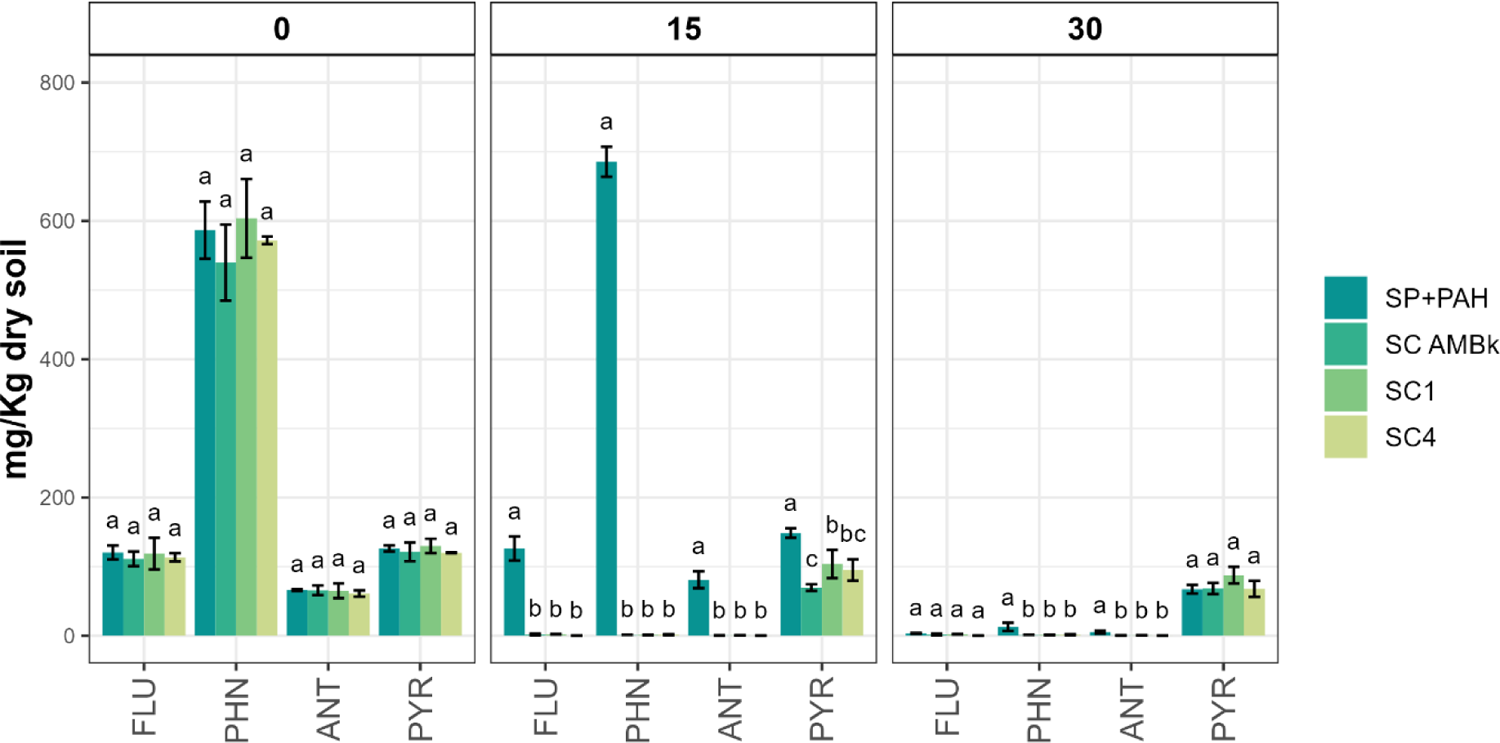
Concentration of supplemented PAHs expressed as mg/Kg of dry soil on day 0 (A) and day 30 (B) of incubation in the inoculated microcosms and in the contaminated control (SP+PAH). The means of independent triplicates with their respective standard deviations are shown. Fluorene (FLU), phenanthrene (PHN), anthracene (ANT) and pyrene (PYR). The non-contaminated control (control) and day 58 of all treatments were excluded from figure, as the concentration of PAHs was below the detection limit.

A significant decrease (p<0.05) of the supplemented PAHs was observed on day 15 of incubation in the inoculated microcosms. Fluorene, anthracene and phenanthrene removal was greater than 99%, with no significant differences between these microcosms. Pyrene degradation was significantly higher (p<0.05) in SC AMBk inoculated microcosms (42% degradation was achieved) compared to those inoculated with SC1 (19%). While no degradation of the supplemented PAHs was observed after 15 days of incubation in SP+PAH, a complete elimination of fluorene, 97% degradation of phenanthrene, 92% anthracene and 46% pyrene were measured after 30 days of incubation. At this point, low molecular weight PAH concentration in the inoculated microcosms was under the quantification limit, while in SP+PAH remaining concentrations of 12.5 mg. Kg-1 of dry soil for phenanthrene and 5.14 mg. Kg-1 of dry soil for anthracene were detected. Concentration of all PAH was below the detection limit in all the studied microcosms at the end of the incubation period (day 58).

### 3.2 Population dynamics and quantification of specific genes related with PAH degradation by qPCR

The dynamics and the degrading potential of soil bacterial populations in the community were assessed by qPCR of the 16S rRNA gene and of two specific genes related: the PAH-RHDα GN gene, and the C230 gene.

The copy number of the 16S rRNA gene for all microcosms during the incubation time was around 10^9^-10^10^ copies.g^-1^ of dry soil. In order to normalize the variations in the copy number of specific genes, a percentage ratio was calculated using the copy number value of each specific gene compared to the one of 16S rRNA gene. Figure 2 shows these gene ratios over time for the different microcosms. No differential pattern was observed in the C230 gene/16S rRNA gene ratio during the incubation period or between microcosms. However, in the inoculated microcosms, the PAH-RHDα GN/16S rRNA ratio was higher than the control and SP+PAH on day 0. After 30 days of incubation this ratio increased its value in SP+PAH reaching similar values to the ones obtained for inoculated microcosms. At the end of the incubation period the PAH-RHDα GN/16S rRNA ratio was similar for all microcosms.

**Figure 2:**
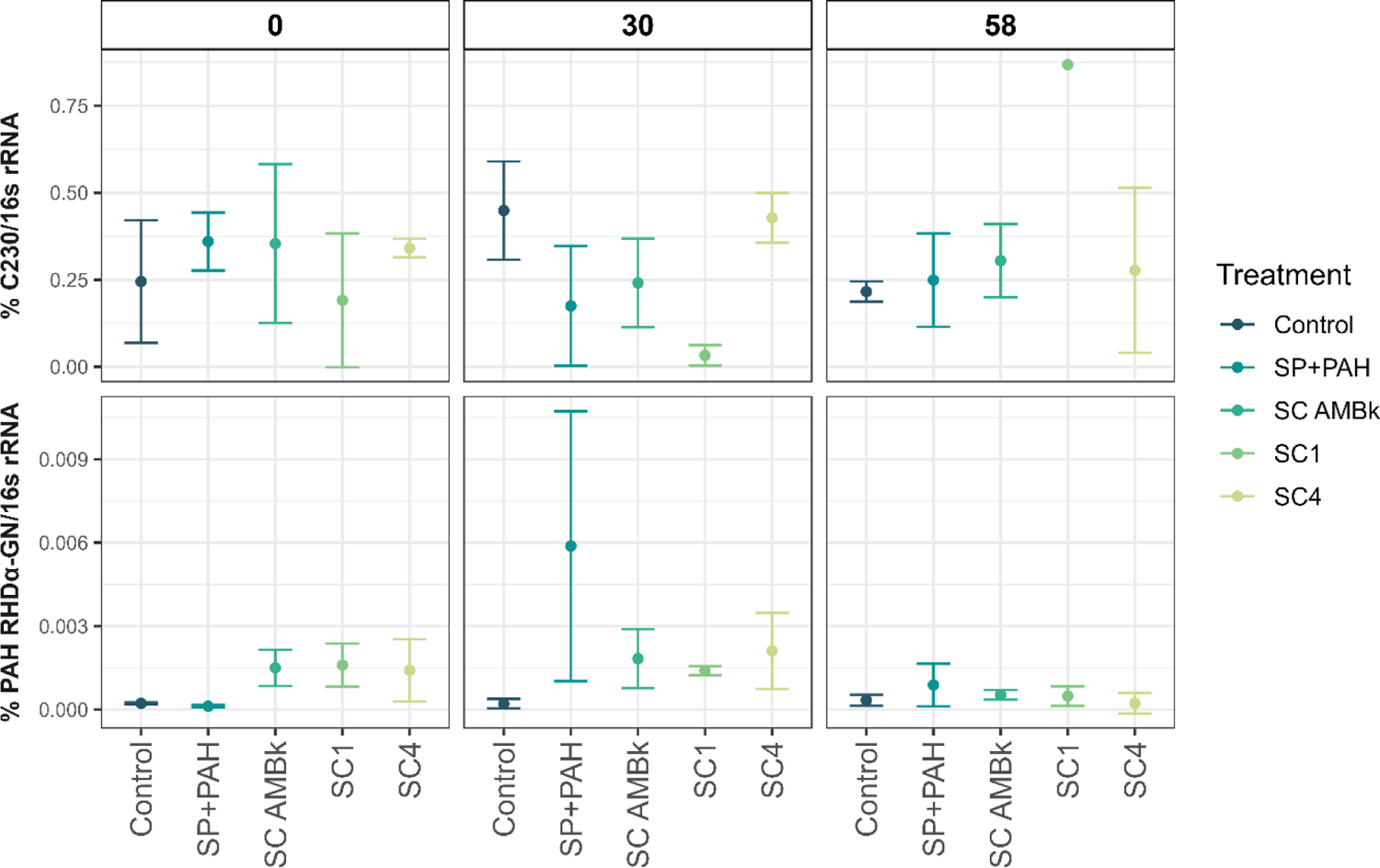
Percentage ratio between the number of copies of the specific gene (C230 gene or PAH RDHα-GN gene) and the number of copies of the 16S rRNA gene for the inoculated microcosms and the controls. The means of triplicate are plotted with their respective standard deviations.

### 3.3 Analysis of the soil microbiome diversity

The inoculation effect on bacterial community structure during the bioaugmentation process was studied through 16S rRNA gene metabarcoding at different times of incubation. Time points were selected considering the observed PAH elimination kinetics (day 15 and 58 for all treatments and day 0 only for the control).

Alpha diversity analysis was carried out applying Chao1, Shannon and Faith indexes (Figure S2). The control microcosms showed an increase in these index values on day 15, being significantly higher than the inoculated microcosms. At the end of the incubation period, no differences were observed among all systems.

Beta diversity analysis showed that inoculation had a remarkable effect on community structure during the first days of incubation (Figure S3), while a convergence in the community structure was observed between inoculated and control microcosms at the end of incubation period.

### 3.4 Composition of the soil microbiomes and monitoring of the inoculated strains

Five predominant phyla were identified in the microcosms on day 15 of incubation: Proteobacteria (32%), Actinobacteriota (15%) Acidobacteriota (7%) and Verrucomicrobiota (6%). Table S1 and Figure 3 show the relative abundance of the bacterial orders at the different incubation times. After 15 days, Galelialles order (8%, Actinobacteriota phylum) was predominant in the control microcosms, followed by the Nitrososphaerales order (6%, Crenarchaeota phylum). In all contaminated microcosms, an increase in Proteobacteria was observed compared to day 0. While its relative abundance in the inoculated microcosmos was 50%, in SP+PAH and control microcosms was 37% and 21% respectively. Within this phylum, Burkholderiales was the predominant order in the microcosms inoculated with SC AMBk, SC4 and in SP+PAH. At genus level, while *Burkholderia-Caballeronia-Paraburkholderia* were the most abundant genera in the SC AMBk and SC4 inoculated microcosms, reaching a relative abundance value of 16% in both systems, it represented only 3.5% in SC1 inoculated microcosms, and did not exceed 1% in the SP+PAH and control microcosms. The ASV identified for Bk (Table S2) contributed to more than 90% to these genera in SC AMBk and SC4 inoculated microcosms, while represented less than 50% in the other microcosms. Within the mentioned order, *Cupriavidus* was a predominant genus in SP+PAH microcosms (15%), reaching relative abundances lower than 4% in the inoculated microcosms, while it did not exceed 1 % in the control microcosms.

**Figure 3.**
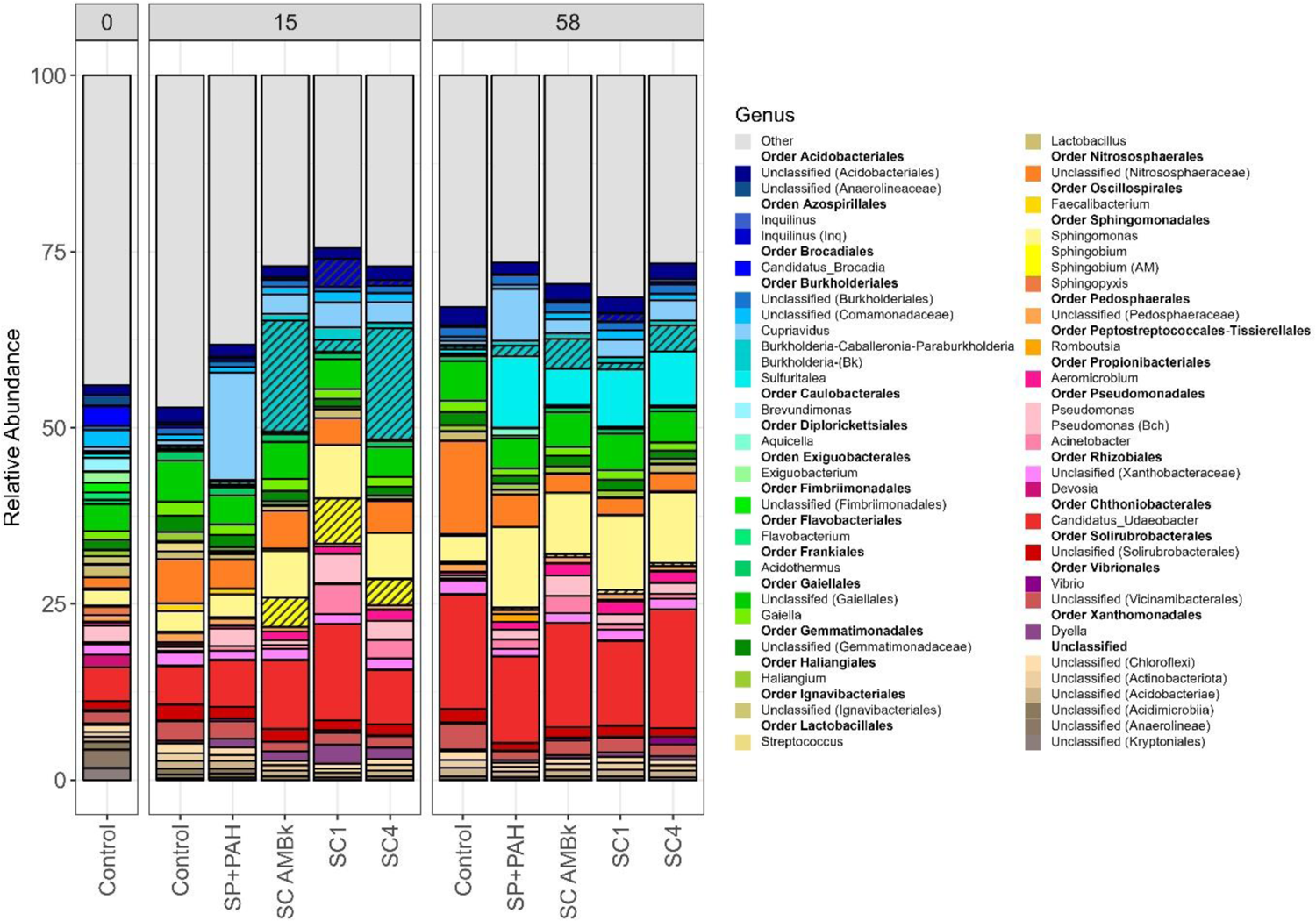
Relative abundances of the genera whose abundance exceeds 1% in the microcosms analyzed at 15 and 58 days of incubation. These genera represented more than 50% of the relative abundance of the community. The average of the triplicates for each treatment is shown. The dashed plot identifies the contribution of ASVs identified with each of the strains to the corresponding genus.

In SC1 inoculated microcosms, Sphingomonadales (Proteobacteria) and Chthoniobacterales (Verrucomicrobiota) were predominant orders, both with relative abundances of 14%. Within the former order, while *Sphingobium* and *Sphingomonas* were major genera in the inoculated microcosms, showing abundances greater than 4% and 5%, respectively, in the SP+PAH microcosms their relative abundances were 1% and 3% respectively. The ASV identified as AM strain (Table S2) was present in almost all microcosms but its contribution to *Sphingobium* genus was more than 99% in the inoculated microcosms. Regarding Chthoniobacterales members, *Candidatus Udeobacter* was the predominant genus (13%) in the SC1 inoculated microcosms and was the second most abundant in the microcosms inoculated with SC4 (9%), SC AMBk (7%) and in the SP+PAH (6%) and control (5%) microcosms.

Among other predominant genera, while *Acinetobacter* (order Pseudomonadales) showed a relative abundance greater than 3% in the SC1 and SC4 inoculated microcosms, in the remaining microcosms it did not exceed values of 0.5%. *Inquilinus* (order Azospirilales) was found in SC1 inoculated microcosms with a relative abundance of 4% at 15 days of incubation, although it did not exceed 1% in the other analyzed microcosms. The ASV identified as Inq strain contributed more than 90% to this genus.

The predominance of Proteobacteria was also observed at day 58 of incubation, in the contaminated microcosms (42-47%), followed by Verrucomicrobiota (13-18%). In contaminated microcosms, *Candidatus Udeobacter* was the predominant genus (12-17%), followed by *Sphingomonas* (9-11%) and *Sulfuritalea* (order Burkholderiales) (5-10%).

Regarding to the control, an increase of the relative abundance of *Candidatus Udeobacter* was observed (16%) compared to day 15, which was the major genus, followed by an unclassified genus from the Nitrososphaeraceae family (13%).

### 3.5 Analysis of Compositions of Microbiomes with Bias Correction (ANCOM-BC)

A differential abundance analysis on bacterial genera was performed using the ANCOM-BC methodology (Lin and Peddada, 2020). This methodology considers the compositional nature of the sequencing data, uses absolute abundances, estimates the sample fraction, and performs a theoretical correction for the bias associated with this fraction between samples. In order to obtain robust results, the analysis focused on those genera that exceed a prevalence of 10% (Nearing et al., 2022). Figure 4 shows the differentially abundant genera in the analyzed microcosms at 15 and 58 days of incubation. After 15 days of incubation, 38 bacterial genera resulted differentially abundant, representing a minor percentage of the total genus found (7%) (Figure 4).

**Figure 4:**
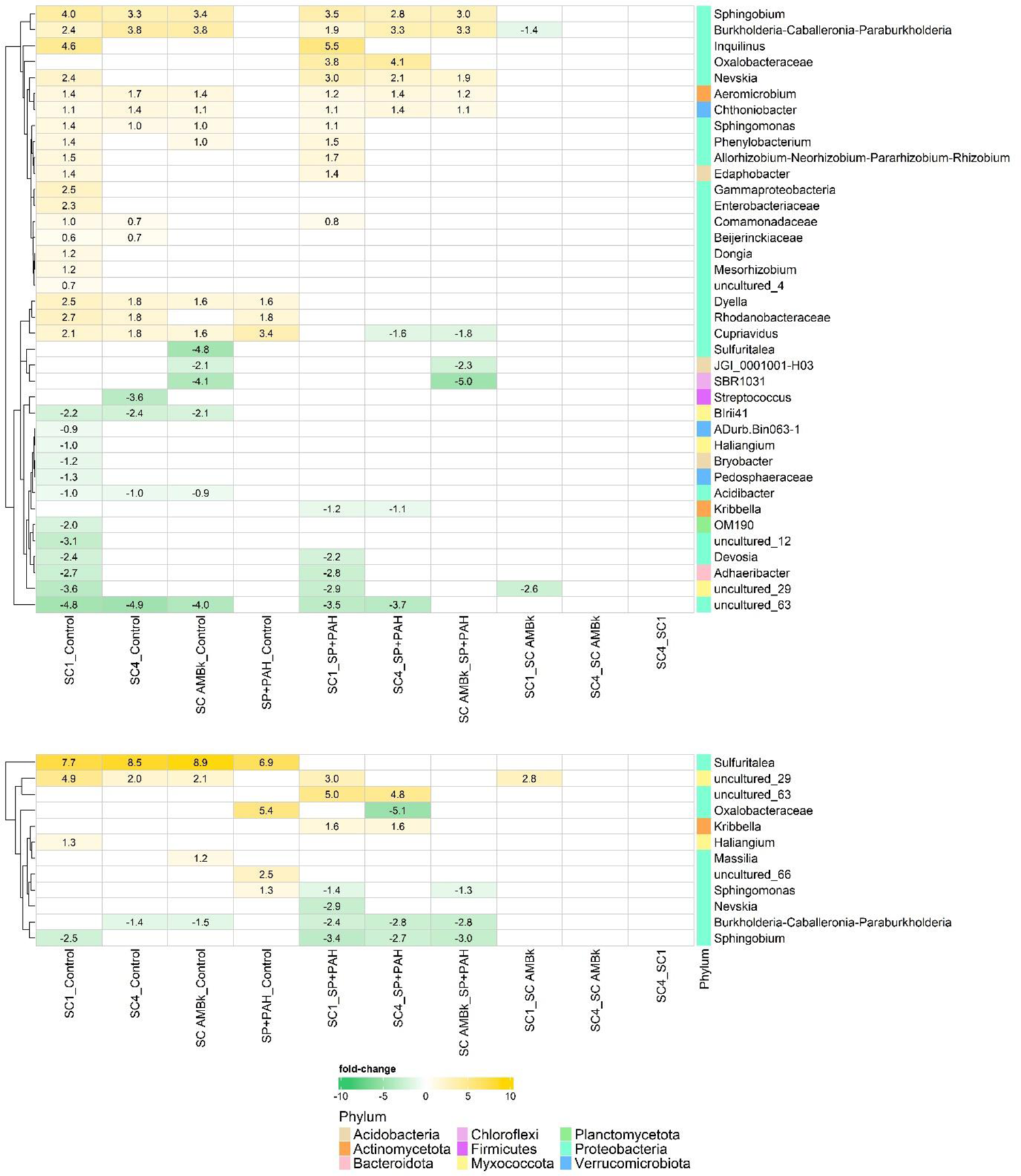
Microbiome genera for which significant changes in absolute abundances were observed between the microcosms analyzed at 15 and 58 days of incubation. Results were expressed as the log fold-change (lfc) of the comparisons between treatments.

When comparing SP+PAH with control microcosms, 3 genera showed significantly higher abundances in the former: *Cupriavidus* (lfc 3.8, Burkholderiales), a genus from the *Rhodanobacteracea* family and *Dyllela* (lfc 1.8 and lfc 1.6 respectively, Xanthomonadales). However, higher differences were observed between the inoculated and the control microcosms. SC1 inoculated microcosms showed the highest number of differentially abundant genera, 21 genera with higher abundance than the control. Among these, major changes were observed for *Inquilinus* (lfc 4.6, Azospirillales), *Sphingobium* (lfc 4.0, Sphingomonadales), and an unclassified genus of the family Rhodobactereaceae (lfc 2.7, Rhodobacterales). 12 genera showed lower abundances in SC1 than in the control, and among them were *Adhaeribacter* (lfc −2.7, Cytophagales), *Devosia* (lfc −2.4, Hyphomicrobiales) and *BIrii41* (lfc −2.2, Polyangiales).

SC AMBk and SC4 inoculated microcosms showed 14 differentially more abundant genera compared to the control, with 10 of them showing the same behavior. Two of the most abundant genera were *Sphingobium* and *Burkholderia-Caballeronia-Paraburkholderia* (Bukholderiales) (lfc 3.3-3.4 and lfc 3.8 respectively). When focusing on the genera being differentially less abundant in SC AMBk and SC4 compared to the control, a non-cultivable genus of the Microscillaceae family (lfc −4.9) and *Streptococcus* (lfc −3.6, Lactobacillales) was found for the SC4 comparison and *Sulfiritalea* (lfc −4.8, Burkholderiales) for the SC AMBk one.

Seven genera were found to be significantly more abundant in the 3 inoculated microcosm compared to the control, *Sphingobium* and *Sphingomonas* (Sphingomonadales), *Aeromicrobium* (Propionibacteriales), *Chthoniobacter* (Chthoniobacterales), *Burkholderia—Caballeronia-Paraburkholderia* and *Cupriavidus* (Burkholderiales) and *Dyella* (Xanthomonadales), while 3 genera showed lower abundances in these microcosms, *Birii41* (Polyangiales), *Acidibacter* and a non-cultivable genus from the Microscillaceae family.

When comparing the inoculated microcosms with SP+PAH, the SC1, SC4 and SC AMBk inoculated microcosms showed 17, 9 and 8 differentially abundant genera, respectively. Although most of these genera behaved similarly to what was observed in the comparison between inoculated and control microcosms, some genera showed unique variation in relation to SP+PAH. For example, a higher abundance of a genus of the Oxalobacteracea family (Burkholderiales) was observed in SC1 (lfc 3.8) and in SC4 (lfc 4.1) inoculated microcosms compared to SP+PAH. Among the genera that showed significantly lower abundances in relation to SP+PAH, *Cupriavidus* was found in the microcosms with SC AMBk (lfc −1.6) and SC4 (lfc −1.8) and *Kribbella* (Propionobacterales) was found in the microcosms with SC1 (lfc −1.2) and SC4 (lfc −1.1).

When comparing the different inoculants, only *Burkholderia-Caballeronia-Paraburkholderia* (lfc −1.1) and a non-cultivable genus of the Sandaracinaceae family (lfc −3.6) showed significantly lower abundances in the SC1 compared SC AMBk inoculated microcosms. No differentially abundant genera were observed when comparing SC4 microcosms with SC1 or SC AMBk microcosms.

At 58 days of incubation, 2% of the total bacterial genera (12 genera) showed significant changes in the comparisons (Figure 4). Only 3 differentially abundant genera were found between the inoculated microcosms and the control, being one of them *Sulfuritalea* (lfc 7.7, 8.5 and 8.9 for SC1, SC4 and SC AMBk respectively). SP+PAH microcosms showed a total of 4 significantly more abundant genera compared to the control: *Sulfuritalea* (lfc 6.9), *Sphingomonas* (lfc 1.3), an unclassified genus of the Oxalobactereaceae family (lfc 5.4) and a non-cultivable genus of the Moraxellaceae family (lfc 2.5).

In relation to the inoculated microcosms, SP+PAH showed significantly higher abundances *Burkholderia-Caballeronia-Paraburkholderia* and *Sphingobium* genera at 58 days of incubation.

### 3.6 Functional prediction by PICRUST and contribution of the different genera to the KOs related to PAH degradation

The functional potential of the bacterial community of each treatment was predicted using PICRUSt2 software. The reliability of this prediction was evaluated by analyzing NSTI values, resulting in an average of 0.21±0.03 for all treatments (Table S3). A total of 7605 KOs were found in the dataset, but only 167 were related to the degradation of aromatic compounds. In order to evaluate which of these KO were differentially abundant in each treatment along the incubation time, ANCOM-BC method was applied performing different comparisons. The Figure S4 shows the comparison between treatments after 15 and 58 days of incubation which resulted in 73 and 19 differentially abundant KO, respectively.

Regarding the comparison at day 15 with SP+PAH, 16 KO were found to be differentially more abundant in SC1 and 10 KO in SC4 and SC AMBk. Seven of these KO were common to the inoculated microcosms and were related to the first steps of the PAH degradation pathway (K14581) and the ortho-cleavage of central intermediates (K00455, K04100, K04101, K05921 and K10219). Furthermore, several KO were found to be significantly more abundant in SP+PAH than in the inoculated microcosms; 15 of these KO were shared between the three inoculated microcosms and were related to toluene (K00055, K15760, K15761, K15762, K15763, K15764, K15765), benzoate (K07535, K07536, K10621, K10622) and phthalate (K18068) degradation.

In the comparisons within the inoculated microcosms, only 3 KO resulted significantly less abundant in SC AMBk than in SC1, but no difference was found between SC4 and SC1 or SC AMBk.

At day 58 of incubation, only one KO (K14581) was significantly more abundant in SP+PAH than in the non-contaminated control and no differences were found in the comparison including the inoculants. The genera contribution to the eight more differentially abundant KO shared among the inoculated microcosm at day 15 were evaluated considering their participation in upper and the lower degradation pathway (Figure 5). For this analysis only the genera contributing in more than 5% in at least one condition were selected. Notably, in the chord diagrams of the inoculated microcosms (Figure 5) *Sphingobium* and *Burkholderia* were the genera that contributed more to the eight analyzed KO. Those related to protocatechuate degradation were mostly assigned to *Sphingobium* (K10219, K04100, K04101 and K11949) and *Burkholderia* represented a minor fraction in the contribution. However, the ones related to 3,4-dihydroxyphenylacetate and benzoate degradation were assigned to *Burkholderia* (K14581, K00455, K05921 and K00141) and no contribution of *Sphingobium* was found in those KO. *Pseudomonas*, *Cupriavidus*, *Acidothermus*, *Reynella* and *Bradirhizobium* genera were also predicted to contribute to the analyzed KO and their contribution was higher in SC1 than in the other treatments (Figure 5).

**Figure 5:**
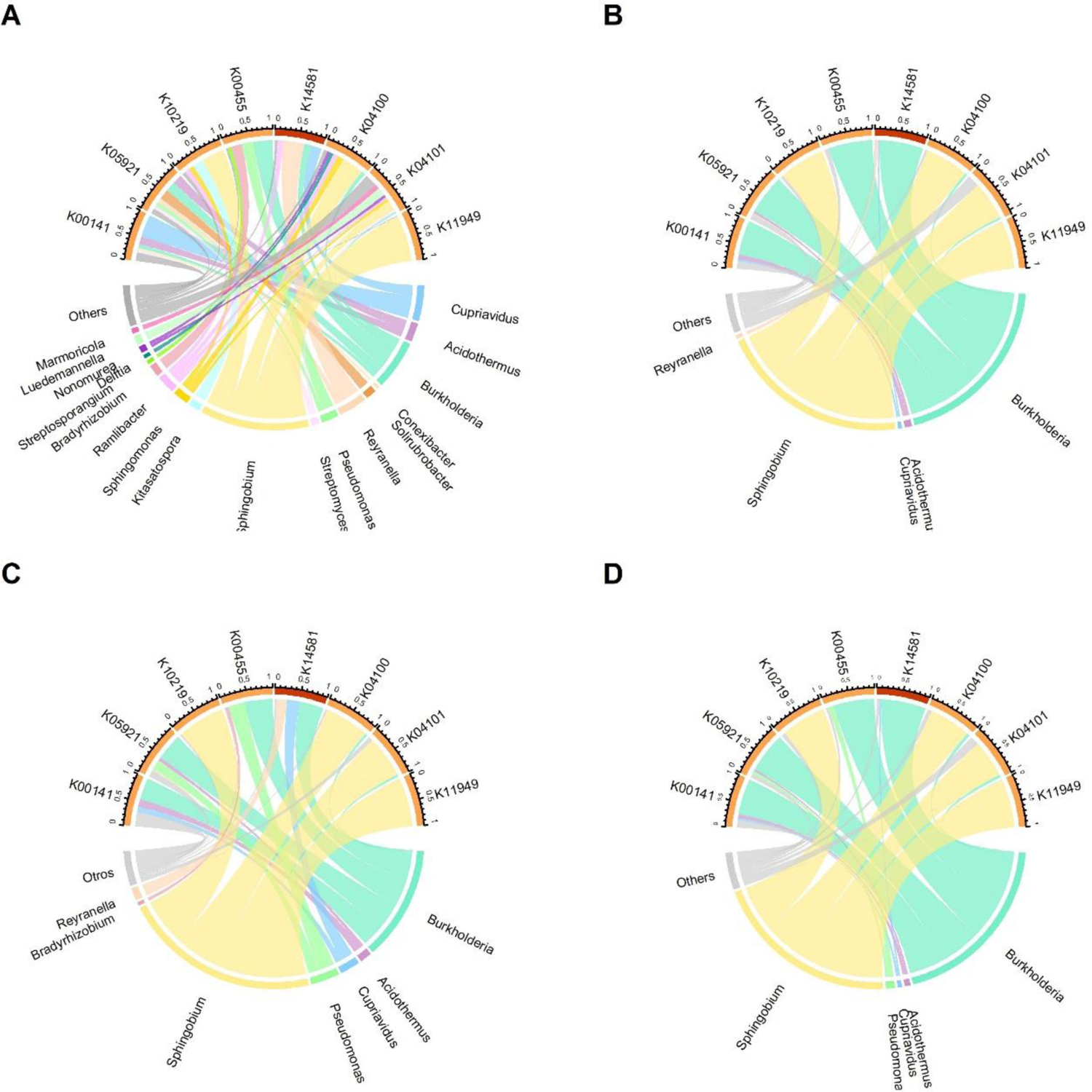
Chord diagram showing the relative contribution for 5 KOs predicted by PICRUSt2, which showed significant differences in SP+PAH contaminated microcosms (A), inoculated with SC AMBk (B), SC1 (C) and SC4 (D) at 15 days of incubation. The averages of the relative contributions are shown for the genera whose relative contributions were greater than 5% in at least one route. Those KOs related to the lower PAH degradation pathway are shown in orange while the KO linked to the high degradation pathway is shown in red.

### 3.7 Soil microbiomes co-occurrence networks

A network analysis was adopted to explore the effects of addition of contaminants and bioaugmentation within soil bacterial interactions. Properties of each resultant network are summarized in Table 1. At day 15 of incubation a decrease in the number of the predicted interactions was observed within the soil bacterial community members due to PAH supplementation, since SP+PAH, as regards to the non-contaminated control, presented a reduction of 38% (from 725 to 450) of total interactions. This reduction was accompanied by a change in the structural features, nodes and edges of bacterial networks as the path length increased and the cluster coefficient and network density decreased, showing that it was a less connected community with more alienated relationships, compared to the control (Table 1 and Figure 6).

**Figure 6:**
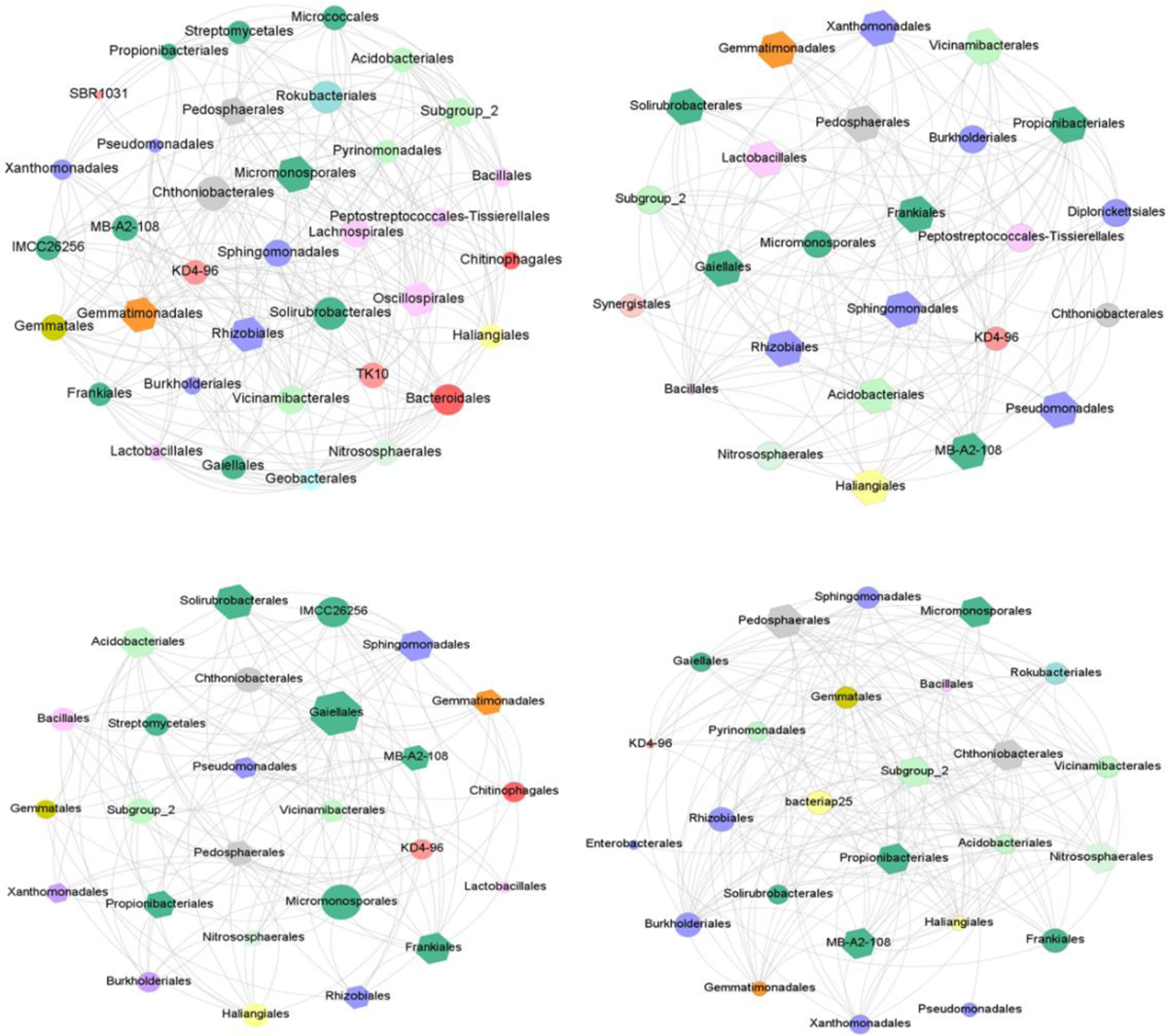
Network co-occurrence analysis of the microbiomes in SP+PAH (A) and inoculated (B) microcosms at day 15 of incubation. A connection stands for a strong (SparCC r > 0.5) and positive correlation (p value < 0.05). The size of each node is proportional to the number of connections (degree). Each node is colored based on the bacterial phyla and keystone taxa are indicated as hexagons.

**Table 1:**
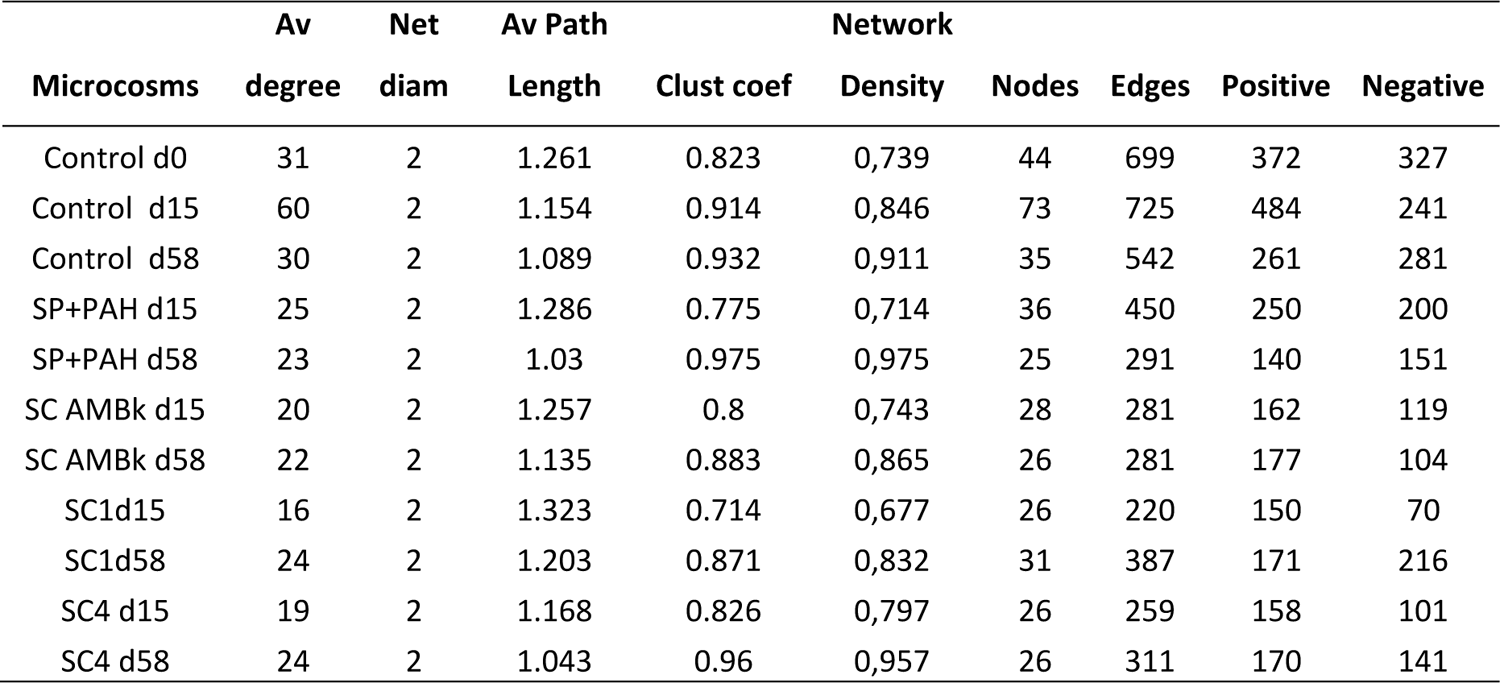
Network properties of soil bacterial communities in analyzed microcosms.

When comparing the inoculated microcosms with the SP+PAH microcosms, on day 15, higher number of interactions was observed in the latter, but this decrease is mainly due to the decrease in negative interactions (around 40 to 70 %).

In all inoculated microcosms, positive interactions were predicted between the orders of the inoculated microorganisms and Propionibacteriales. Also, positive interactions were found in SC1 and SC4 inoculated microcosms with Xanthomonadales (Figure 6).

Regarding the co-occurrence networks of the inoculated microcosms, SC AMBk and SC4 resultant networks possessed the shortest average path length, the highest cluster coefficient and network density (Table 1), indicating closer relationships of soil microbial taxa.

At the end of the incubation time, all the co-occurrence networks had a higher density value, higher clustering coefficient and shorter path length value compared to those of the day 15, indicating that the nodes had a higher degree of interaction and association when the contaminant concentration and inoculants members decreased.

The number of orders identified as keystone varied between treatments and incubation time. However, common keystone taxa were found in the contaminated and inoculated microcosms, such as Gaiellales, Gemmantimonadales, Actinomycetales, Solirubrobacterales and Sphigomonadales,.

In addition, to study the interactions existing prior to inoculation, a network for SP+PAH microcosms at genus level was constructed (Figure S5). In this network interactions between soil bacterial residents and *Burkholderia-Caballeronia-Paraburkholderia* were analyzed. As a result, negative interactions were found with *Cupriavidus, Dyella* and *Sulfuritalea* genera and positive interactions with *Aeromicrobium, Gaiella* and *Sphingomonas* genera.

## 4. Discussion

With an increasing focus on microbiome engineering, understanding and manipulating complex microbial ecosystems has become a defining mission for microbiome science (Kong et al., 2018). Our study provides insights into the structure, dynamics, and ecology of interacting microbial species during inoculation of allochthonous microorganisms capable of removing contaminants, which is a promising strategy in bioremediation of acutely contaminated soils (Nwankwegu et al., 2022).

The bioaugmentation with three consortia used in this work stimulated the degradation of the mixture of PAHs supplemented to the soil, reaching the complete elimination of low molecular weight PAHs and partial degradation of high molecular weight PAH already at 15 days of incubation (Figure 1). The higher biodegradation potential of PAH (number of copies of the PAH RHDα-GN), observed in the inoculated microcosms compared to SP+PAH and control microcosms (Figure 2), was expected due to the introduction of the inoculants that contain those genes in their genomes (Macchi et al., 2021).

These results were reinforced by the functional prediction carried out by PICRUSt2 (Figure 4 and 5), where the inoculated genera *Sphingobium* and *Burkholderia* contributed more to the KOs that showed higher abundance than SP+PAH and control microcosms, highlighting their main role on PAH degradation among the consortium members. Within these KOs, K14581 (naphthalene 1,2-dioxygenase ferredoxin reductase component) could act in the first steps of the degradation of PAHs (fluorene, anthracene, benzo[a]pyrene and pyrene) (Zhou et al., 2020) with *Burkholderia* genus contributing the most. Expression of this enzyme by Bk was observed during the degradation of phenanthrene in liquid media, both in SC-4 and in SC AMBk (Macchi et al., 2021; Nieto et al., 2023). This could be related to the higher pyrene degradation found in SC AMBk inoculated microcosms at 15 days of incubation (Figure 1). Several KOs linked to the degradation of protocatechuate (K04100, K04101, K10219, K00455), a metabolite of PAH degradation, showed higher abundances in the inoculated microcosms, which could indicate the predominance of the ortho-cleavage pathway (Figure S4). Other authors also reported an increase in specific functions associated with inocula in bioaugmentation studies (Cayetano et al., 2021; Zhou et al., 2022).

Microbial community in SP+PAH microcosms were also able to degrade the supplemented PAH; however, this process, unlike the inoculated microcosms, was delayed until day 30 (Figure 1). This could be the result of selecting native degrading microorganisms made by pollutants, consistent with the increase of PAH RHDα-GN gene after 30 days of incubation. No differences were observed in C230 gene copy number between the inoculated microcosms and the control microcosms, probably due to the participation of this gene in other metabolic pathways involving one-ring aromatic compounds.

Bioaugmentation failure could result from consortia selected under laboratory conditions becoming less competitive under harsh and/or variable field conditions (Palkova, 2004; Steensels et al., 2019). Hence, following the survival of the inoculum is key to understand the success of bioaugmentation process. *Sphingobium* and *Burkholderia* were both members of SC AMBk and SC4 consortia and were found to be among the predominant genera in the microcosms inoculated with these consortia at 15 days of incubation. A significant increase of *Burkholderia* abundance was also observed in the SC1 inoculated microcosms in comparison to SP+PAH, even though this genus is not a member of this consortium; this behavior would indicate mutualistic interactions between *Sphingobium* and *Burkholderia*, which were also previously reported (Zhang et al., 2013; Willsey and Wargo, 2015; Festa et al., 2016a).

A recent metanalysis showed that inoculant diversity correlates significantly and positively with inoculum effect (Liu et al., 2023). One explanation could be that more species in a consortium increase its multifunctionality and stability (Jiménez et al., 2018). However, our results did not show differences between the different inoculants, demonstrating that a more diverse consortia will not always lead to an improvement in the degradation performance. In our study, *Sphingobium* and *Burkholderia* played a main role in PAH-degradation, which might be related to the changes in resource availability due to the PAHs addition to the soil. Thus, *Sphingobium* and *Burkholderia* may have taken advantage of this open niche, outcompeting the native community and increasing their establishment success. (Silverstein et al., 2023).

On the other hand, the rest of consortium members are known to play a major role in the lower PAH-degradation pathway (Macchi et al., 2021), whose enzymes are more widespread in nature than the ones in the higher PAH-degradation pathway and could act in other several metabolic pathways. Therefore, the absence of establishment of the other consortium members could be a consequence of their competition with the high diverse native community (Figure S2 and 3), since the remaining niche available, related to PAH intermediate degradation, decreased (Mallon et al., 2015). In this scenario with high resource availability, bioaugmentation efficiency relayed on those consortium members with key catabolic functionalities (i.e., *Sphingobium* and *Burkholderia*), while further degradation of the intermediates, which involve fewer specific functions, could be carried out by the native microorganism. The ability of *Sphingobium* AM as a sole strain to establish and increase the soil biodegradation activity was previously observed (Festa et la., 2016a) when applied as inoculant in an acutely contaminated soil. However, in that study the inoculation produced drastic changes in the structure and diversity of soil bacterial community that remained even after 63 days of incubation.

Moreover, it becomes essential to study the interaction between the inoculants and the native community, since the efficiency of the process depends on the survival of the inoculant but, at the same time, the least impact is sought on the structure and functionality of the native community (Mawarda et al., 2020) after the bioremediation process is complete.

Incubation conditions had a stimulating effect on the bacterial populations of the control microcosms, evidenced by the increase in both richness and diversity at 15 days of incubation. This effect, previously observed in other bioaugmentation processes, was not found in the inoculated microcosms (Figure S2), which could be mainly linked to the predominance of the inoculants (Festa et al., 2016b; Chaudhary et al., 2021; Zhou et al., 2022). In turn, Faith’s index showed a significant decrease in the inoculated microcosms regarding the control and SP+PAH microcosms, probably indicating that inoculation generated a rapid change in the bacterial community towards microorganisms whose phylogenetic relationship was greater. The β diversity analysis also showed an early impact of inoculation on the bacterial community structure (Figure S3).

Co-occurrence networks of the inoculated microcosms (Figure 6) at day 15 showed a lower number of total interactions than in SP+PAH, possibly due to a reorganization of interactions caused by the inocula. Inoculation resulted in a higher decrease in negative than in positive interactions (Table 1) which correlates with bioaugmentation effectiveness, since negative interactions of an inoculant with other soil microorganisms can produce a delay in the effect of bioaugmentation (Jiang et al., 2022).

Changes in community structure were observed as a consequence of both inoculation and contamination. As previously discussed, a selection of PAH-degrading microorganisms was observed in SP+PAH microcosms. As in the inoculated microcosms, Proteobacteria was the main phylum, with a predominance of Burkholderiales order. While the predominant genus in SP+PAH microcosms was *Cupriavidus*, in SC AMBk and SC4 inoculated microcosms its abundance was significantly lower compared to SP+PAH (Figure 3). This could be a consequence of the predominance of *Burkholderia* in inoculated microcosms, since both genera have a similar catabolic potential (Pérez-Pantoja et al., 2012) and could compete for resources, where the predominance of one could limit the growth of the other. Competition between the genera *Cupriavidus* and *Burkholderia* was inferred in the correlation networks of the contaminated system (Figure S5).

Common effects on the soil bacterial community were observed beyond the behavior of the inoculated genera. A predominance of the Proteobacteria phylum (Table S1) was found, which includes several PAH-degrading members. The genera *Aeromicrobium* (Propionobacteriales) and *Nevskia* (Xanthomonadales) showed higher abundances in the inoculated microcosms compared to SP+PAH and control microcosms. Members of these genera were reported to have direct participation in PAH degradation (Crampon et al., 2018; Li et al., 2021). In addition, positive interactions were predicted between the inoculated orders and members of the autochthonous orders potentially involved in PAH degradation such as Propionibacteriales and Xanthomonadales order (Figure 6). Indeed, all the bacterial orders that were indicated as keystone in most of the treatments, meaning they play a key role in modulating network structure and function, were orders with known PAHs-degrading capabilities (Pagé et al., 2015; Zhang et al., 2022).

The structural and functional divergence between the microbiome of the control and the contaminated microcosms decreased at 58 days of incubation with no significant differences observed in diversity indices (Figure S2) and reduced number of differentially abundant genera and functions (Figure 3 and S4). At this time, the increase in network connections and microbial interactions was observed (Table 1), which could indicate a more stable microbial community as more connected networks could have higher community stability and resistance to disturbances (de Vries et al., 2018). When referring to the observed differentially abundant genera at 58 days of incubation, *Sulfuritalea*, a genus found in PAH-contaminated soils (Guo et al., 2017; Yang et al., 2022), was significantly higher in the contaminated microcosms than in the control microcosms (Figure 3 and 4). This genus was pointed as indicator taxa, tolerant to hydrocarbon, potentially involved in hydrocarbon-degradation or benefiting from the degradation by-products (Cébron et al., 2022). Its presence in the contaminated microcosms at day 58 could be linked to a residual effect of the contamination. However, in all microcosms, a predominance of *Candidatus Udaeobacter* was found at the end of the incubation period, one of the most abundant soil bacteria with global dissemination (Brewer et al., 2016; Willms et al., 2020), whose abundance correlates with soil health (Wilhelm et al., 2023).

Results obtained in this work show that the new microbiome assembly achieved through bioaugmentation with the three degrading consortia not only increases the metabolic capacities of the microbiome due to the added microorganisms, but also stimulates the native degrading populations of the soil. At the same time, at the end of the incubation period, the impact of the inoculation was reduced as all the microbiomes converged with the non-contaminated control microcosms (Figure 3, 4 and 5; Table S1).

## 5. Conclusion

The three inoculated consortia were successful in accelerate the removal of contaminants. As long as the inoculant has the capacity to initiate the PAH degradation, competitiveness and efficiency of the inoculant does not depend on its diversity in cases of acutely contaminated soils with PAHs of up to four rings. Furthermore, an increase in the abundance of bacterial genera related to soil health was observed at the end of the of the incubation period. Our results show a significant advantage of inoculation with allochthonous microorganisms since temporary establishment of the inoculant and its low impact on the native microbiota are desired objectives in inoculation processes.

