## Supplemental material for "Challenging the impact of consortium diversity on bioaugmentation efficiency and native bacterial community structure in a freshly PAH-contaminated soil"

**Figure S1:** Rarefaction curves of the analyzed samples, showing the number of ASVs observed in relation to the sequencing depth. The dotted red line indicates the number of sequences used as a cut-off point for rarefaction of the samples.

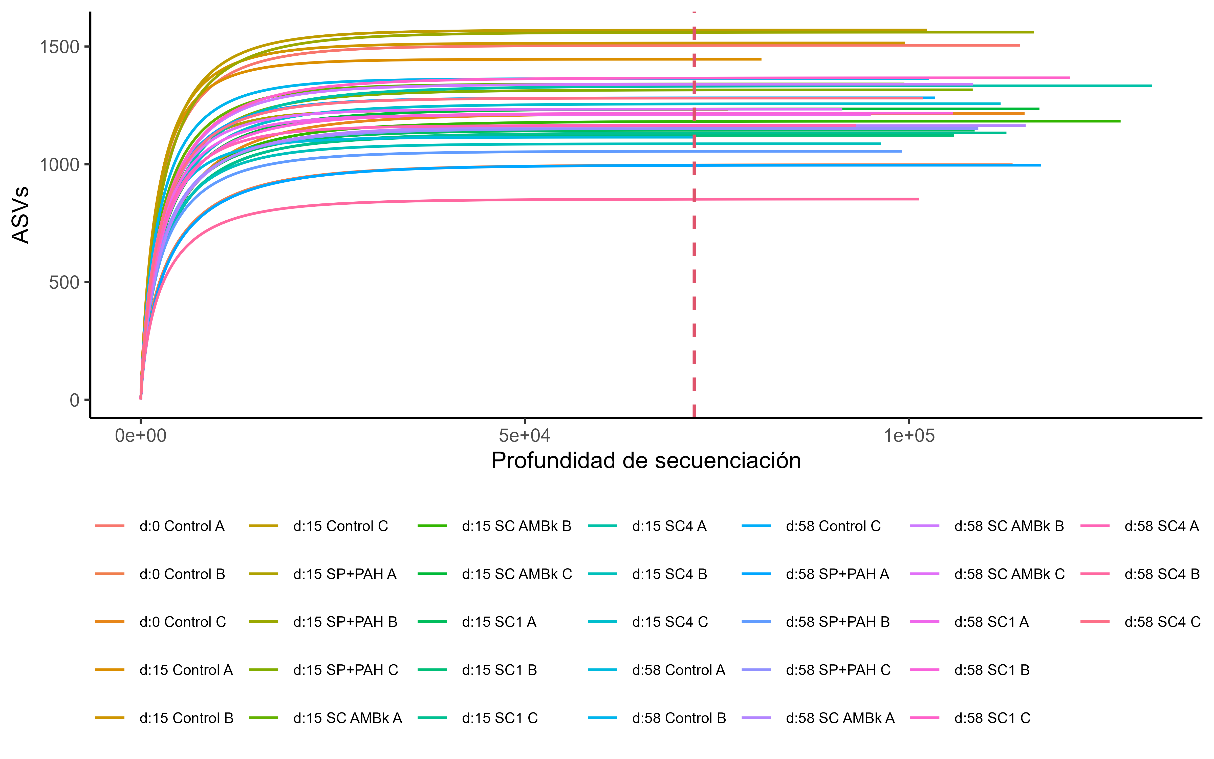

**Figure S2**: Alpha diversity. Chao1, Shannon and Faith indices at 0 (Control), 15 and 58 days of incubation of the analyzed microcosms. The results of triplicates of each system are shown.

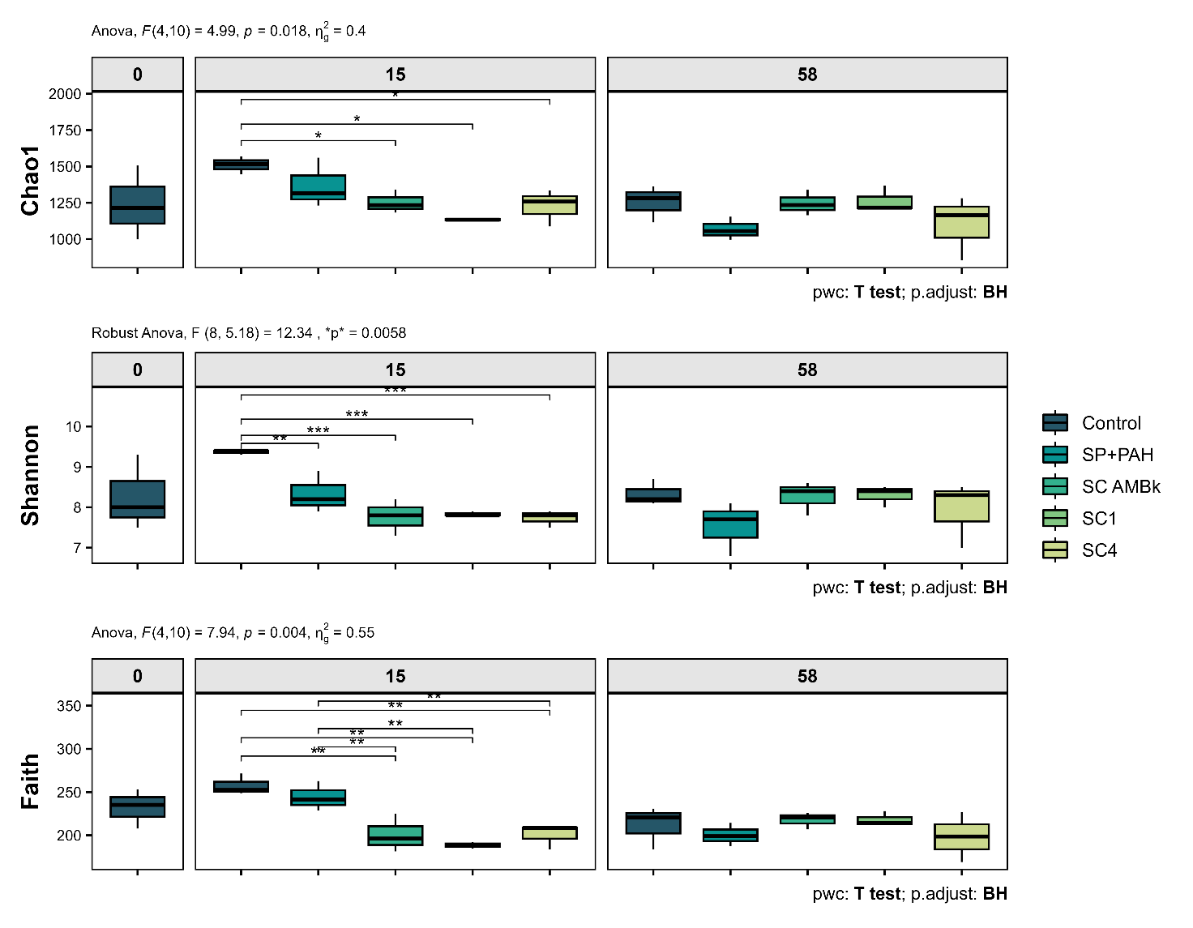

**Figure S3**: Principal component analysis (PCoA) using the Bray-Curtis method for the comparison of the analyzed microcosms at the different incubation times. The two main axes that explain most of the variability are shown. The size of the dots indicates the incubation time, while the colors represent the different treatments.

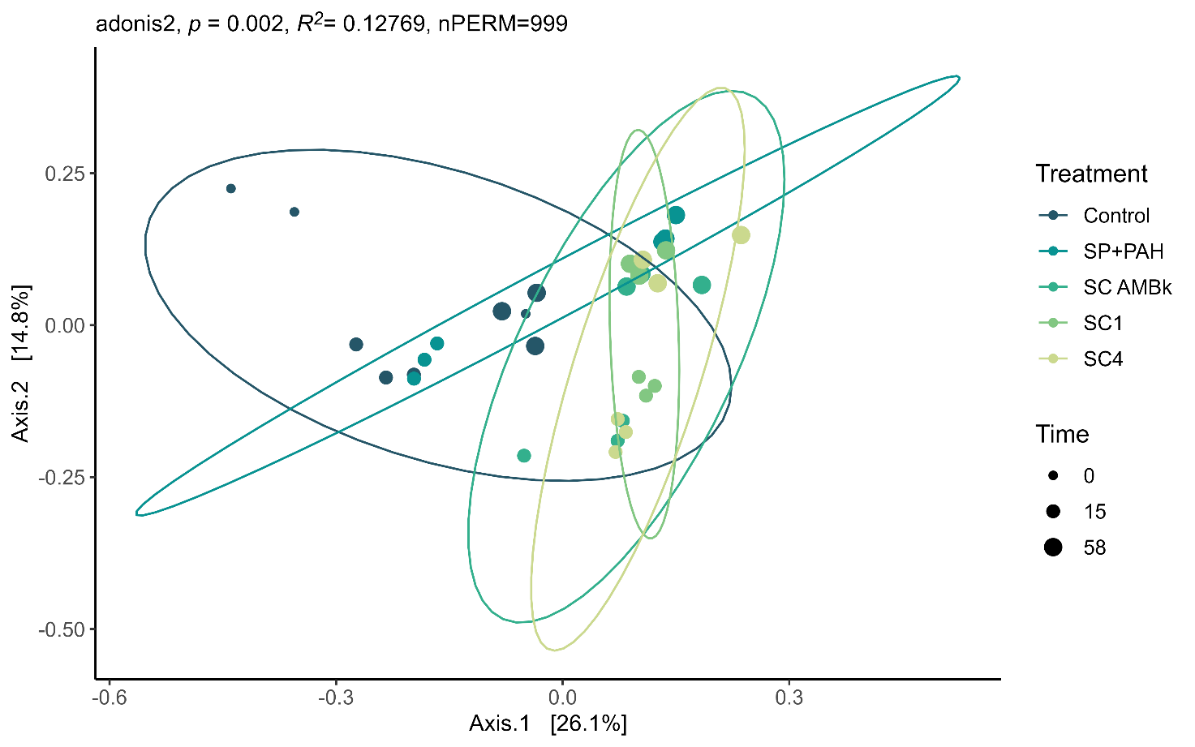

**Table S1**: Relative abundance of the main orders (relative abundance higher than 1% in at least one condition)

| Order | control | control | SP+PAH | SC AMBk | SC1 | SC4 | control | SP+PAH | SC AMBk | SC1 | SC4 |
| --- | --- | --- | --- | --- | --- | --- | --- | --- | --- | --- | --- |
| Burkholderiales | 9,97 | 5,91 | 20,32 | 23,46 | 11,29 | 24,13 | 6,05 | 24,42 | 17,58 | 17,54 | 19,74 |
| Sphingomonadales | 3,63 | 3,14 | 3,46 | 10,85 | 14,04 | 10,25 | 3,98 | 11,88 | 9,17 | 11,18 | 10,47 |
| Chthoniobacterales | 4,85 | 5,6 | 6,71 | 10,1 | 14,04 | 8,2 | 16,46 | 12,46 | 15,12 | 12,41 | 17,26 |
| Gaiellales | 5,2 | 8,15 | 5,82 | 7,3 | 5,93 | 5,85 | 7,5 | 5,4 | 6,46 | 6,73 | 5,78 |
| Nitrososphaerales | 1,63 | 6,3 | 4,16 | 5,42 | 3,84 | 4,57 | 13,24 | 4,58 | 2,67 | 2,46 | 2,65 |
| Rhizobiales | 7,26 | 5,54 | 4,51 | 4,73 | 4,27 | 4,88 | 4,92 | 2,91 | 3,95 | 4,57 | 4,05 |
| Solirubrobacterales | 2,16 | 3,59 | 2,64 | 2,71 | 2,09 | 2,29 | 2,76 | 1,52 | 2,06 | 2,2 | 1,8 |
| Xanthomonadales | 1,3 | 1,63 | 2,21 | 2,15 | 3,9 | 2,77 | 1,58 | 1,22 | 1,64 | 1,8 | 1,54 |
| Acidobacteriales | 1,72 | 2,63 | 2,07 | 2,11 | 2,04 | 2,67 | 3,14 | 2,16 | 2,99 | 3,15 | 2,94 |
| Propionibacteriales | 1,16 | 1,45 | 1,7 | 1,88 | 1,43 | 2,02 | 0,73 | 1,34 | 1,95 | 2,11 | 1,85 |
| Vicinamibacterales | 2,09 | 3,65 | 3,32 | 1,83 | 2,42 | 2,43 | 4,72 | 1,77 | 2,84 | 2,81 | 2,41 |
| Gemmatimonadales | 1,77 | 2,85 | 2,12 | 1,81 | 1,43 | 1,69 | 2,55 | 1,64 | 2,01 | 2,09 | 1,63 |
| Frankiales | 0,82 | 1,74 | 1,47 | 1,45 | 1,14 | 1,17 | 0,96 | 0,62 | 0,83 | 0,89 | 0,7 |
| Pseudomonadales | 2,69 | 0,76 | 3,25 | 1,32 | 8,86 | 5,57 | 0,83 | 2,85 | 5,58 | 2,24 | 2,62 |
| Micromonosporales | 0,73 | 1,43 | 1,28 | 1,19 | 0,98 | 1,09 | 0,96 | 0,71 | 0,78 | 0,91 | 0,67 |
| Pedosphaerales | 1,58 | 2,21 | 1,59 | 0,94 | 0,6 | 0,99 | 1,92 | 0,85 | 1,33 | 1,4 | 1,11 |
| Lactobacillales | 2,17 | 2,89 | 1,24 | 0,87 | 1,27 | 0,31 | 1,38 | 1,51 | 0,18 | 0,07 | 1,27 |
| MB-A2-108 | 0,6 | 1,14 | 0,89 | 0,77 | 0,64 | 0,82 | 1,04 | 0,62 | 0,86 | 0,98 | 0,75 |
| Bacillales | 1,41 | 1,27 | 1,15 | 0,76 | 0,85 | 0,71 | 0,8 | 0,62 | 0,78 | 0,81 | 0,79 |
| Subgroup_2 (Acidobacteriae) | 0,79 | 1,07 | 1,1 | 0,76 | 0,59 | 0,84 | 1,24 | 0,79 | 0,93 | 0,95 | 0,92 |
| KD4-96 | 1,07 | 1,4 | 1,07 | 0,64 | 0,7 | 0,85 | 1,27 | 0,6 | 0,78 | 0,9 | 0,71 |
| Gemmatales | 0,58 | 1,22 | 0,92 | 0,61 | 0,72 | 0,81 | 0,93 | 0,38 | 0,84 | 0,73 | 0,72 |
| Haliangiales | 0,92 | 1,33 | 0,56 | 0,57 | 0,33 | 0,46 | 0,84 | 0,86 | 0,99 | 1,01 | 0,73 |
| IMCC26256 (Acidimicrobiia) | 1,09 | 0,83 | 0,69 | 0,55 | 0,4 | 0,48 | 0,5 | 0,41 | 0,47 | 0,46 | 0,41 |
| Dongiales | 0,19 | 0,33 | 0,35 | 0,49 | 0,75 | 0,58 | 0,2 | 0,35 | 0,36 | 0,41 | 0,31 |
| Caulobacterales | 2,47 | 0,31 | 0,22 | 0,47 | 0,47 | 0,44 | 0,2 | 0,35 | 0,36 | 0,36 | 0,3 |
| Lachnospirales | 0,9 | 1,73 | 1,28 | 0,4 | 0 | 0 | 1,35 | 0,47 | 0,21 | 0,09 | 1,08 |
| Azospirillales | 0,29 | 0,14 | 0,1 | 0,38 | 4,06 | 0,89 | 0,21 | 0,27 | 0,39 | 1,29 | 0,45 |
| Oscillospirales | 0,35 | 1,15 | 1,01 | 0,32 | 0 | 0 | 0,51 | 0,02 | 0,08 | 0,03 | 0,32 |
| Flavobacteriales | 1,02 | 0,05 | 0,12 | 0,29 | 0,04 | 0,08 | 0,05 | 0,02 | 0,02 | 0,02 | 0,02 |
| Polyangiales | 0,69 | 1,17 | 0,38 | 0,21 | 0,12 | 0,15 | 0,64 | 0,31 | 0,39 | 0,52 | 0,3 |
| Peptostreptococcales-Tissierellales | 0,32 | 1,45 | 0,63 | 0,13 | 0,05 | 0,12 | 0,08 | 2,38 | 0,12 | 1,29 | 0,14 |
| Sphingobacteriales | 1,32 | 0,04 | 0,18 | 0,09 | 0,03 | 0,09 | 0,01 | 0 | 0,01 | 0,01 | 0,01 |
| Vibrionales | 0,25 | 0,22 | 0,43 | 0,07 | 0,3 | 0,13 | 0,16 | 0,13 | 0,47 | 0,83 | 1,08 |
| Diplorickettsiales | 0,12 | 0,05 | 0,04 | 0,05 | 0,09 | 0,07 | 0,4 | 1,11 | 0,31 | 0,32 | 0,24 |
| Anaerolineales | 1,67 | 0,47 | 0,31 | 0,03 | 0 | 0 | 0,01 | 0 | 0,01 | 0,01 | 0 |
| Exiguobacterales | 1,56 | 0 | 0 | 0 | 0 | 0,01 | 0 | 0 | 0 | 0 | 0 |
| SBR1031 | 2,79 | 0,54 | 0,6 | 0 | 0,01 | 0,01 | 0,02 | 0,01 | 0,02 | 0,02 | 0 |
| Brocadiales | 2,79 | 0,36 | 0,42 | 0 | 0 | 0 | 0 | 0 | 0 | 0 | 0 |
| Ignavibacteriales | 1,8 | 0,26 | 0,29 | 0 | 0 | 0 | 0 | 0 | 0 | 0 | 0 |
| Kryptoniales | 1,67 | 0,25 | 0,3 | 0 | 0 | 0 | 0 | 0 | 0 | 0 | 0 |
| Fimbriimonadales | 1,44 | 0,17 | 0,21 | 0 | 0 | 0 | 0 | 0 | 0 | 0 | 0 |
| Others | 21,17 | 23,58 | 18,88 | 12,29 | 10,28 | 11,58 | 15,86 | 12,46 | 14,46 | 14,4 | 12,23 |

**Table S2**: ASVs identified in the microcosm samples analyzed that showed the highest percentage of identity for each of the inoculated strains, obtained from BLASTn. The 16S RNA gene sequence of the T strain was the only one to which no ASV could be assigned, since the ASV identified with the greatest similarity did not reach an identity percentage greater than 99%.

| Cepas | Nombre del ASV identificado | % de identidad |
| --- | --- | --- |
| *Burkholderia sp.* Bk | 86f1248ee4ea85afd8c5b67cc1b04d4c | 100 |
| *Sphingobium sp.* AM | 99fee6617d20f02a0e0641f4d5a57916 | 99,61 |
| *Inquilinus limosus* Inq | b53094561082f2158e468e437235ead1 | 100 |
| *Pseudomonas sp.* Bc-h | 472466279b75a9efd1ce4fbc71e9f0bd | 100 |
| *Pseudomonas sp.* T | 472466279b75a9efd1ce4fbc71e9f0bd | 97,68 |
| *Klebsiella aerogenes* B | 945184b6386c192c0066e0a98a154780 | 99,61 |

**Table S3:** NSTI values for each sample from PICRUSt2 prediction

| Muestra | NSTI |
| --- | --- |
| d:0 SP Ct A | 0,25 |
| d:0 SP Ct B | 0,24 |
| d:0 SP Ct C | 0,13 |
| d:15 SP Ct A | 0,25 |
| d:15 SP Ct B | 0,25 |
| d:15 SP Ct C | 0,23 |
| d:15 SP+PAH A | 0,22 |
| d:15 SP+PAH B | 0,23 |
| d:15 SP+PAH C | 0,21 |
| d:15 SP SC AMBK A | 0,21 |
| d:15 SP SC AMBK B | 0,19 |
| d:15 SP SC AMBK C | 0,21 |
| d:15 SP SC1 A | 0,19 |
| d:15 SP SC1 B | 0,17 |
| d:15 SP SC1 C | 0,17 |
| d:15 SP SC4 A | 0,18 |
| d:15 SP SC4 B | 0,20 |
| d:15 SP SC4 C | 0,21 |
| d:58 SP Ct A | 0,25 |
| d:58 SP Ct B | 0,25 |
| d:58 SP Ct C | 0,25 |
| d:58 SP+PAH A | 0,20 |
| d:58 SP+PAH B | 0,19 |
| d:58 SP+PAH C | 0,22 |
| d:58 SP SC AMBK A | 0,20 |
| d:58 SP SC AMBK B | 0,23 |
| d:58 SP SC AMBK C | 0,24 |
| d:58 SP SC1 A | 0,22 |
| d:58 SP SC1 B | 0,22 |
| d:58 SP SC1 C | 0,22 |
| d:58 SP SC4 A | 0,23 |
| d:58 SP SC4 B | 0,18 |
| d:58 SP SC4 C | 0,22 |

**Figure S4**: KOs linked to the degradation of aromatic compounds which were differentially abundant in the microcosms analyzed after 15 and 58 days of incubation. Color intensity reflects the magnitude of the logarithm of the fold change (lfc). Each column indicates the compared pair, where secondly the reference condition for the comparison is indicated.

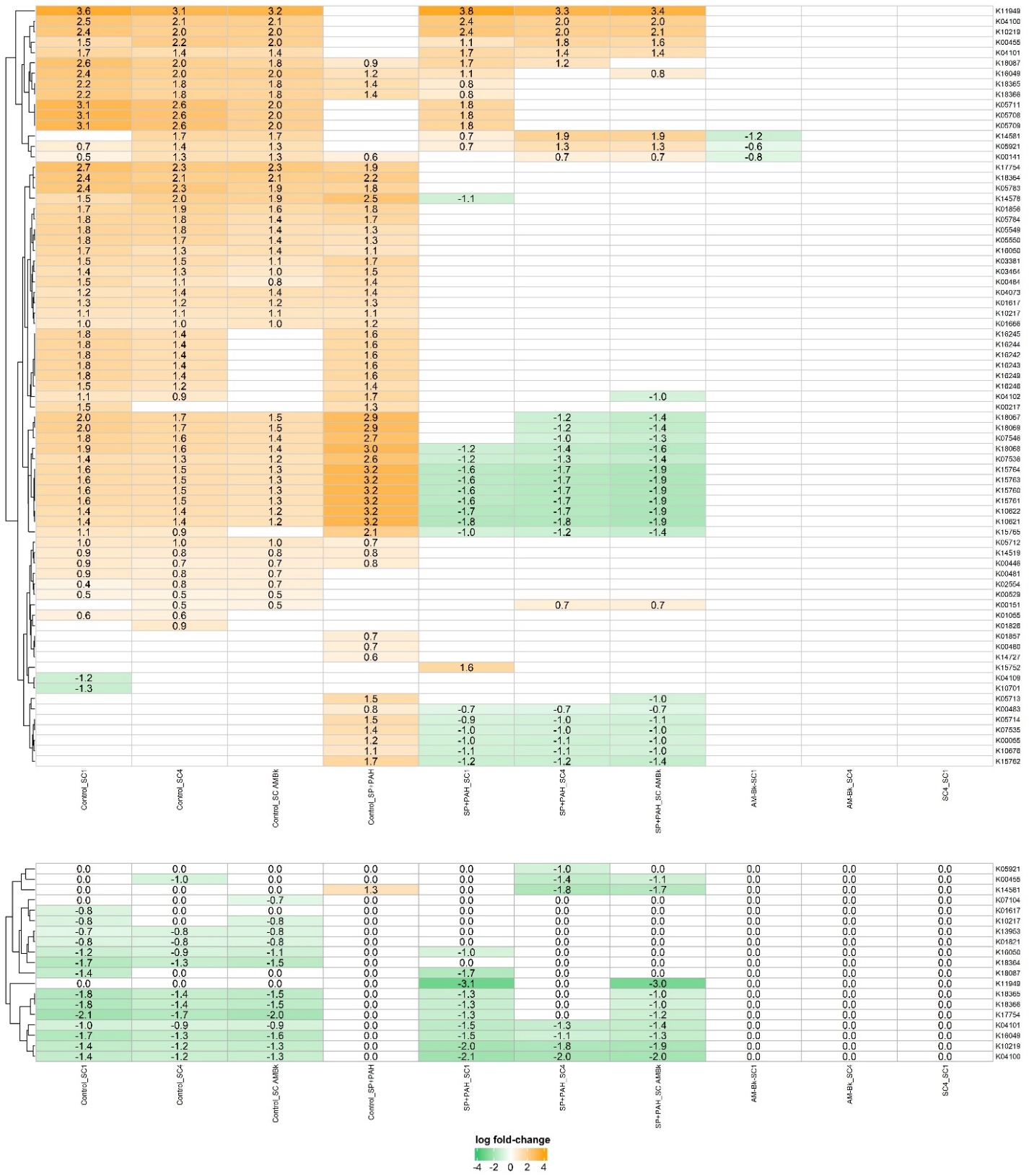

**Figure S5:** Network co-occurrence analysis of the microbiomes of the SP+PAH microcosms at genus level. A connection stands for a strong (SparCC r > 0,5) and significant (p value < 0.05) correlation (Blue: negative. Red: positive). The size of each node is proportional to the number of connections (degree). Each node is colored based on the bacterial phyla.

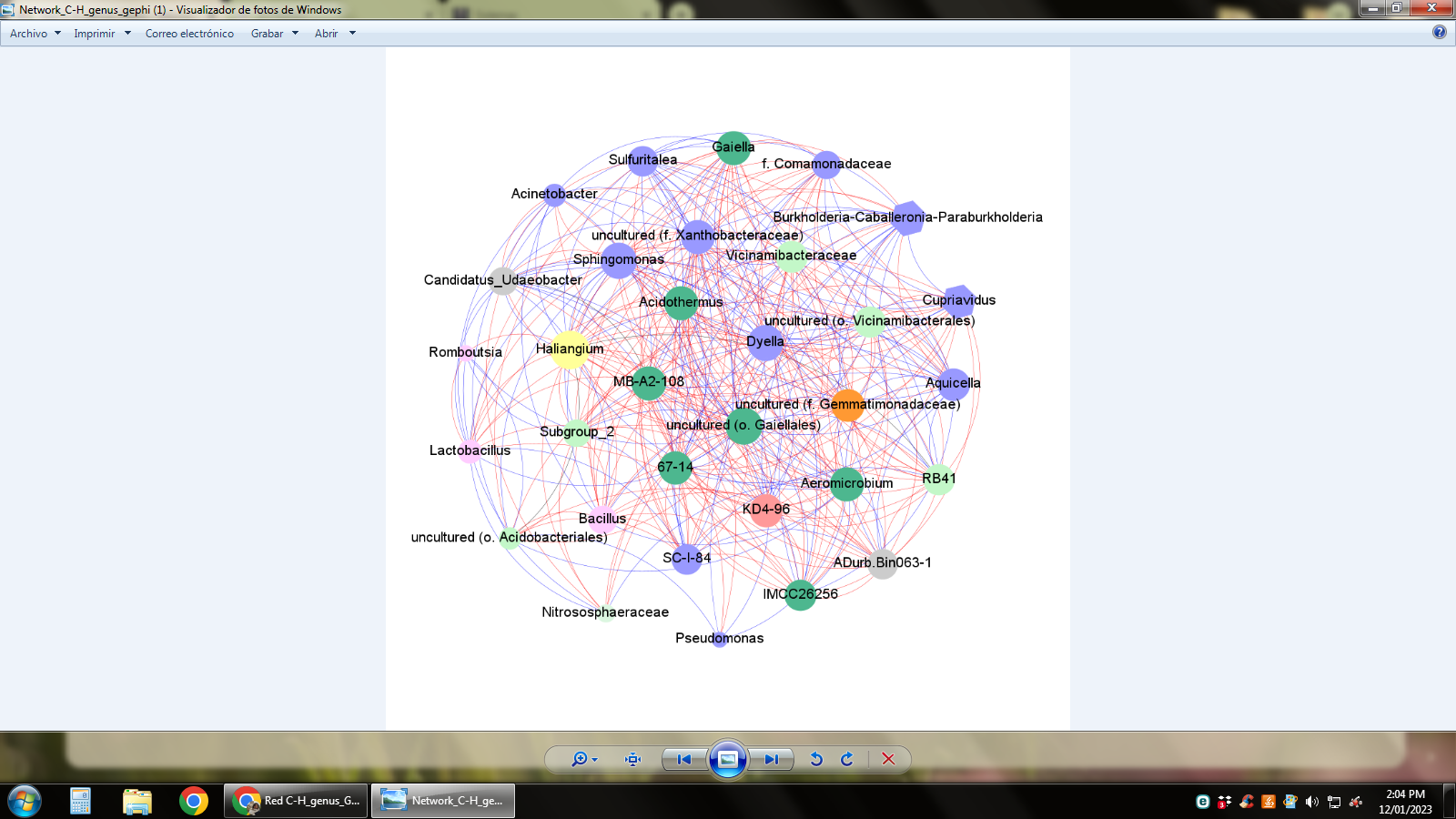
